# Sex specific effects of environmental toxin-derived alpha synuclein on enteric neuronal-epithelial interactions

**DOI:** 10.1101/2024.11.27.625701

**Authors:** Hayley N Templeton, Alexis T Ehrlich, Luke A Schwerdtfeger, Julietta A Sheng, Ronald B Tjalkens, Stuart A Tobet

**Affiliations:** Department of Biomedical Sciences, Colorado State University, Fort Collins, CO, USA; Department of Environmental and Radiological Health Sciences, Colorado State University, Fort Collins, CO, USA; School of Biomedical Engineering, Colorado State University, Fort Collins, CO, USA; Ann Romney Center for Neurological Disease, Harvard Medical School, Brigham and Women’s Hospital, Boston, MA, USA

**Keywords:** Parkinson’s disease, sex differences, alpha synuclein, calcitonin gene related peptide (CGRP), goblet cells

## Abstract

**Background:** Parkinson’s Disease (PD) is a neurodegenerative disorder with prodromal gastrointestinal (GI) issues often emerging decades before motor symptoms. Pathologically, PD can be driven by accumulation of misfolded alpha synuclein (aSyn) protein in the brain and periphery, including the GI tract. Disease epidemiology differs by sex, with men twice as likely to develop PD. Women, however, experience faster disease progression, higher mortality, and more severe GI symptoms. Gut calcitonin gene related peptide (CGRP) is a key regulator of intestinal contractions and visceral pain. The current study tests the hypothesis that sex differences in GI symptomology in PD are the result of aSyn aggregation altering enteric CGRP signaling pathways.

**Methods:** To facilitate peripheral aSyn aggregation, the pesticide rotenone was administered intraperitoneally once daily for two weeks to male and female mice. Mice were sacrificed two weeks after the last rotenone injection and immunohistochemistry was performed on sections of proximal colon.

**Key Results:** Levels of aSyn were heightened in myenteric plexus neurons and a subset of neurons immunoreactive to CGRP in rotenone treated mice. Female mice exhibited 153% more myenteric aSyn, 26% more apical CGRP immunoreactivity, and 66.7% more aSyn in apical CGRP^+^ fibers after rotenone when compared to males. Goblet cell numbers were diminished but the individual cells were larger in the apical regions of crypts in the colons of rotenone treated mice.

**Conclusions:** This study used a mouse model of PD to uncover sex specific alterations in enteric neuronal and epithelial populations, underscoring the importance of considering sex as a biological variable while investigating prodromal GI symptoms.

**Graphical Abstract:** 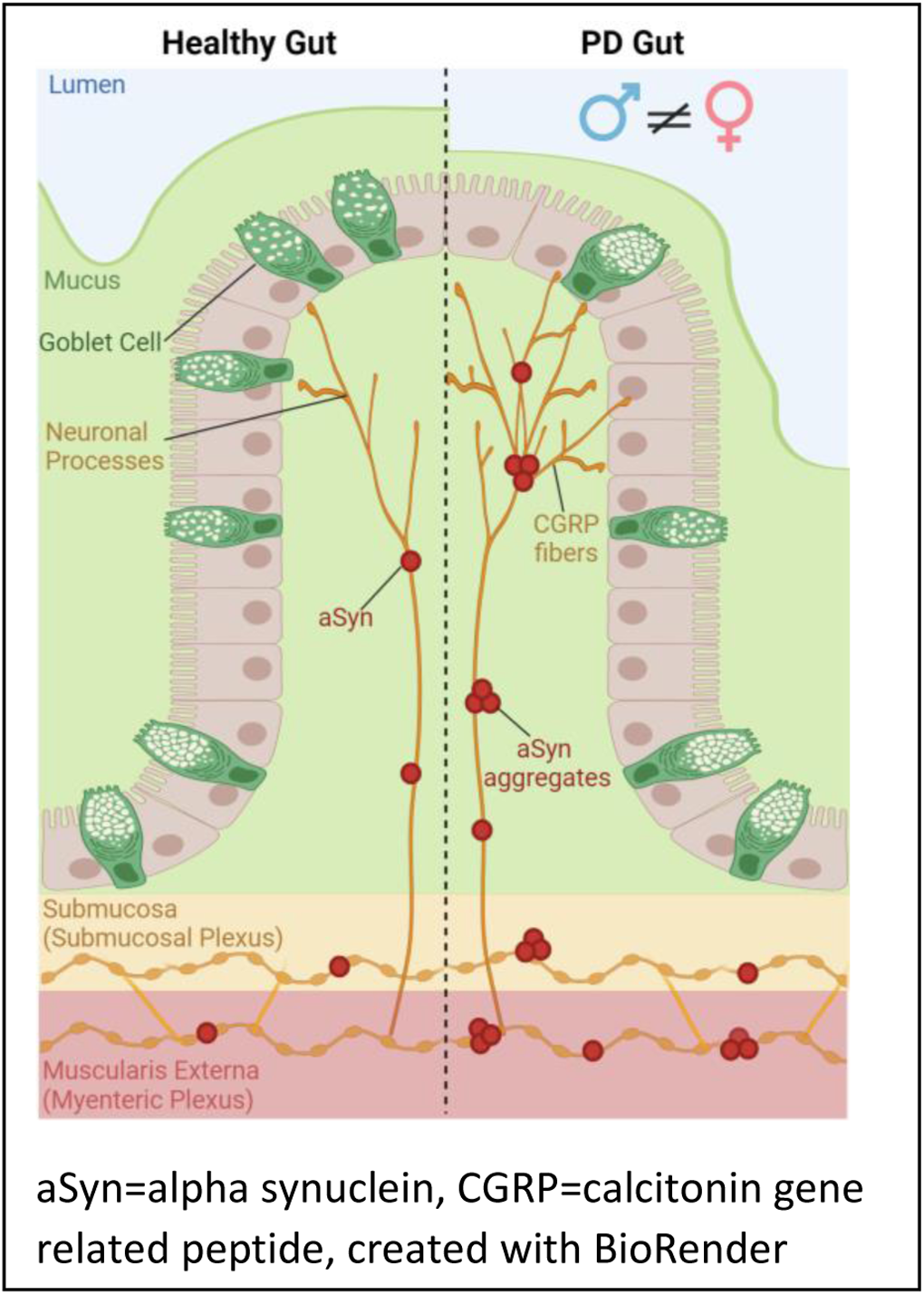

**KEY POINTS:**

- Mouse model of Parkinson’s Disease (PD) was used to investigate sex specific impact of enteric alpha synuclein (aSyn) on colonic goblet cells and CGRP^+^ neurons and fibers.
- Sex specific alterations in intestinal neuronal and epithelial signaling pathways in response to aSyn provides insight into sex differences in PD etiology and prodromal gastrointestinal symptoms.

## 1 INTRODUCTION

Parkinson’s disease (PD) is a neurodegenerative disorder with prodromal symptoms that emerge decades before motor dysfunction. Gastrointestinal (GI) symptoms including bloating, visceral pain, gastroparesis, and constipation, are seen in 10 – 70% of PD subjects^1,2^ and can manifest up to 20 years before diagnosis.^3^ GI symptoms impact quality of life, and pose significant health risks including, malnutrition, intestinal obstruction, and impaired medication effectiveness.^4,5^ PD is driven pathologically by the accumulation of misfolded alpha synuclein (aSyn) protein in the brain and periphery, including the GI tract. Environmental toxins such as rotenone exacerbate aSyn aggregation, producing PD-like pathology and symptoms when administered to rodents.^6^ Gut pathology in rotenone PD models includes peripheral gastroparesis^7^ and decreased intestinal transit,^8,9^ creating a viable murine model for investigating mechanisms of GI dysfunction in PD.

Females with PD exhibit faster disease progression and higher mortality rates than males^10^. In addition, females with PD report more GI symptoms with higher severity,^11–13^ including constipation and abdominal pain, symptoms that heavily impact quality of life^14^ and are among the earliest PD symptoms.^15,16^ Aggregation of aSyn in the intestinal wall alters gut immune populations^17^, and sex differences in immune responses are abundant systemically,^18^ and peripherally in the gut mucosal immune system.^19^ Roles of peripheral immune influence on enteric neurons and epithelial populations involved in prodromal PD gut symptomology are not well elucidated. Evidence on the role of sex in neurodegenerative disease^11,20,21^ along with the current study underscores the importance of including sex as a biological variable in gut focused PD investigations.

Calcitonin gene related peptide (CGRP) is a neuropeptide implicated in regulation of GI motility, inflammation, and nociception through dorsal root ganglia (DRG) afferents^22^ and intrinsic enteric neurons.^23,24^ There are two forms of CGRP, α and β, which share around 90% homology^25^ and have nearly identical biological activity.^26,27^ While not extensively characterized, there are likely classes of intrinsic sensory neurons (IPANs) in the myenteric plexus that express βCGRP^28,29^ and αCGRP^30^. Therapeutic interventions targeting αCGRP significantly inhibit colonic transit, leading to constipation,^31,32^ while elevated αCGRP is associated with diarrhea suggesting that αCGRP plays a role in GI motility.^31^ Gut mucus regulates intestinal transit by acting as a lubricant to decrease physical resistance, and altered mucus levels influence GI transit time.^33,34^ CGRP produced by enteric neurons modulates goblet cell mucus secretion,^35^ as well as motility^36^, further pointing towards CGRP involvement in intestinal transit regulation. Accumulation of aSyn in enteric neurons likely influences gut motility, driven by cholinergic neurons.^37,38^ However, the impact of aSyn accumulation in subsets of enteric cholinergic neurons, including CGRP^+^ cells, on mucosal neuroepithelial signaling remains to be seen. Determining if enteric CGRP^+^ neurons and mucosal projecting fibers accumulate aSyn in a mouse model of PD would provide valuable data towards understanding the altered GI function observed in PD.

To investigate the relationship of sex in aSyn accumulation in enteric CGRP fibers and provide a read out of goblet cell morphological alterations, a rotenone model of PD was utilized. Levels of aSyn were quantified in CGRP expressing myenteric neurons and mucosal fibers. In addition, goblet cell anatomic localization and morphology were analyzed. Data from the present study underscores the importance of examining sex selective mechanisms in PD related GI dysfunction, providing crucial insights into some of the earliest prodromal symptoms in PD.

## 2 MATERIALS AND METHODS

### 2.1 Animals

Male and female C57BL/6J background mice aged 3-4 months were used for these studies. All animal protocols were approved by the Institutional Animal Care and Use Committee (IACUC) and Colorado State University Lab Animal Resources under protocol CoA # 1657. Mice were housed on a 12-h light/dark cycle with access to standard chow (Envigo TD.2918) and water *ad libitum*.

### 2.2 Rotenone Preparation and Injections

Rotenone was prepared as a 50x stock dissolved in 100% dimethyl sulfoxide (DMSO) and stored in amber septa vials at −20°C. This solution was then diluted in medium-chain triglyceride, Miglyol 812 N, to obtain a final concentration of 2.5 mg/mL rotenone in 98% Miglyol 812 N, 2% DMSO. Rotenone was prepared fresh every other day. Mice were weighed every day and received intraperitoneal injections at a dose of 2µL/g body weight once daily for 14 days. The optimal dose was determined in a previous publication.^39^ Fourteen days after the last rotenone injection, mice were deeply anesthetized with isoflurane and terminated via decapitation. The intestine was removed from stomach to colon and placed in 4% paraformaldehyde (PFA) at 4°C for 24 h. Tissue was subsequently placed in 1X phosphate buffered saline (PBS) at 4°C until sectioning.

### 2.3 Immunohistochemistry

50 µm thick sections were prepared from 1-3 mm pieces of proximal colon and submerged in 4% agarose. The tissue was in agarose for a total of 9 minutes: 5 min on a room temperature shaker, and 4 min at 4°C to ensure polymerization. Tissue was cut at a thickness of 50 µm on a vibrating microtome (VT1000S; Leica microsystems, Wetzlar, Germany). Sections were washed in 1x PBS for 10 minutes at 4°C and incubated in 0.1M glycine for 30 min at 4°C, followed by three 5 min PBS washes. Next, tissue sections were incubated in 0.5% sodium borohydride at 4°C for 15 min, followed by three 5 min PBS washes. Sections were blocked in PBS with 5% normal goat serum (NGS; Lampire Biological, Pipersville, PA), 1% hydrogen peroxide, and 0.3% Triton X (Tx) for 1 hr at 4°C with a change of solution at 30 min. Following blocking, sections were placed into primary antisera or lectin outlined in Table 1.

**Table 1.** Primary antibodies used.

| Primary Antibody | Immunizing Antigen | Raised | Dilution | Source | Catalog Number |
| --- | --- | --- | --- | --- | --- |
| PGP9.5 | Recombinant full length human UCHL1 purified from E. coli | Chicken | 1:500 | Novus Biologicals | NB110-58872SS |
| CGRP | Full length rat α-CGRP | Rabbit | 0.625μg/mL | ImmunoStar | 24112 |
| aSyn | Rabbit α-synuclein (D37A6) | Rabbit | 1:500 | Cell Signaling | 4179 |

#### Goblet Cell Lectin Histochemistry

The lectin Ulex Europaeus Agglutinin I conjugated with Rhodamine (UEA-1; Vector Labs) was used at a concentration of 0.625µg/mL in PBS containing 5% NGS and 0.3% Tx for 2 days at 4°C. Sections were then washed four times at 15 min intervals in PBS with 0.02% Tx, followed by four PBS washes. Finally, sections were mounted on slides, and cover slipped with an aqueous mounting medium (Aqua-Poly/Mount, Polysciences, Warrington, PA).

#### PGP9.5 and aSyn Immunohistochemistry

Sections were incubated for 2 days at 4°C in primary antisera containing anti-Protein Gene Product (PGP) 9.5 (Novus Biologicals, Centennial, CO) at 1:500 and anti-alpha-Synuclein (Cell Signaling, Danvers, MA) at 1:500 in PBS containing 5% NGS and 0.3% Tx. The aSyn antibody labels total aSyn per the vendor and several publications.^40–42^ Following primary antisera incubation, sections were washed at room temperature in PBS with 1% NGS and 0.32% Tx four times at 15 min intervals. Sections were then incubated for two hours in secondary antibodies to the appropriate species (Alexa Fluor 405 anti-rabbit and Alexa Fluor 568 anti-chicken; Table 2) at 1:500 constituted in PBS with 0.32% Tx. Following secondary antibody incubation, sections were washed four times at 15 min intervals in PBS with 0.02% Tx, followed by four PBS washes. Finally, sections were mounted on slides, and cover slipped with an aqueous mounting medium (Aqua-Poly/Mount, Polysciences, Warrington, PA).

**Table 2.** Secondary antibodies used.

| Secondary Antibody | Conjugate | Dilution | Source | Catalog Number |
| --- | --- | --- | --- | --- |
| Goat anti-rabbit IgG | Alexa Fluor 405 | 1:500 | Invitrogen | A-31556 |
| Goat anti-chicken IgY | Alexa Fluor 568 | 1:500 | Abcam | ab175477 |
| Goat anti-rabbit IgG | Alexa Fluor 594 | 1:500 | Invitrogen | A-11037 |
| Goat anti-rabbit IgG | Alexa Fluor 647 | 1:500 | Invitrogen | A-31573 |
| Biotin-SP donkey anti-rabbit IGG | Biotin-SP | 1:2500 | Jackson ImmunoResearch | 711065152 |

#### CGRP and aSyn Immunohistochemistry

Sections were incubated for 2 days at 4°C in primary antisera containing anti-aSyn (Cell Signaling, Danvers, MA) at 1:500 in PBS containing 5% normal donkey serum (NDS) and 0.3% Tx. A negative control of the aSyn antibody revealed no signal (**Supplemental Fig. 1C**). Sections were then washed four times at 15 min intervals in PBS with 1% NDS and 0.32% Tx at room temperature. Next, sections were incubated for 2 hr at room temperature in biotinylated donkey-anti-rabbit (Table 2, Jackson ImmunoResearch Laboratories Inc, West Grove, PA) at 1:2500 constituted in PBS with 0.32% Tx. Sections were washed four times at 15 min intervals in PBS with 0.02% Tx at room temperature before incubation in Alexa Fluor 594-streptavidin (Invitrogen) at 1:500 for 1 hr. Sections were then washed four times at 15 min intervals in PBS with 0.02% Tx at room temperature. Sections were incubated for 2 days at 4°C in primary antisera containing anti-αCGRP (ImmunoStar, Hudson, WI) at 0.625µg/mL in PBS containing 5% NGS and 0.3% Tx. Of note, this antibody likely labels a small percentage of βCGRP, however, signal was only partially extinguished in βCGRP pre-absorption studies performed by the vendor. A pre-absorption with full length αCGRP peptide (AnaSpec, Fremont, CA) eliminated signal (**Supplemental Fig. 1A**) and there was no signal in controls with secondary antisera omitted (**Supplemental Fig. 1B**). Following primary antibody incubation, sections were then washed four times at 15 min intervals in PBS with 0.02% Tx before incubation in Alexa Fluor 647 (Table 2, Invitrogen, Waltham, MA) at 1:500 for 2 hr at room temperature. Sections were then washed four times at 15 min intervals in PBS with 0.02% Tx, followed by four PBS washes. Finally, sections were mounted on slides, and cover slipped with an aqueous mounting medium (Aqua-Poly/Mount, Polysciences, Warrington, PA).

### 2.4 Tissue Imaging & Analysis

Images were taken using a Zeiss LSM800 upright confocal laser scanning microscope and a 40x or 63x (EC Plan-Neofluar) oil immersion objective. High resolution images were taken using a Zeiss LSM880 upright confocal microscope with an Axiocam 503 monochromatic camera and a 63x (Plan-Apochromat) oil immersion objective. Data for CGRP and aSyn and goblet cell analyses was gathered from confocal Z-stacks with 15 planes 1 µm apart with the top and bottom planes set from the focus of labeling. Max intensity Z-projections were performed, and regions of interest (ROIs) were defined as the apical or basal half of a crypt. Apical and basal regions were defined by measuring the length of a crypt and dividing it into two equal parts. ROIs were gathered from 12 crypts per animal and averaged. aSyn and PGP 9.5 analyses were gathered from individual, single plane confocal images. Control areas for aSyn and PGP9.5 colocalization were initially defined as muscle regions with no PGP9.5-ir. These regions had no aSyn-ir. Twelve 50 µm x 50 µm ROIs containing aSyn and PGP 9.5 double labeling were selected per sample. aSyn and CGRP analyses were gathered from 20 µm z-projection confocal images. The apical half of six crypts and six myenteric neurons containing aSyn and CGRP double labeling were selected per animal from 20 µm z-projection confocal and subsequently averaged. All image acquisition and analyses were performed by a researcher blinded to treatment. All image analyses were performed using Fiji (v1.0; NIH). Goblet cell quantity and size was measured using the analyze particles tool in Fiji. Images were thresholded based on optical density and a binary watershed was applied prior to quantification. Mean goblet cell size was calculated based off of the area as measured by analyze particles for individual anatomic ROIs as described above. Immunoreactive percentage of area labeled was calculated using the ‘measure’ tool in Fiji on thresholded images restricted to the defined ROI for each respective analysis. To determine the % area of aSyn in CGRP neurons/fibers, thresholded CGRP-IR images were used to generate ROIs using the analyze particles tool. Thresholded aSyn-IR images were then analyzed using CGRP ROIs as area restrictions. Then, the percentage of total CGRP-IR area that contained aSyn-IR was calculated.

### 2.5 Statistics

All statistical analyses were performed using Graphpad Prism version 10.1.2 (Graphpad Software, La Jolla, CA). For aSyn and PGP 9.5 average area, CGRP area, and percent area of aSyn in CGRP, a two-way ANOVA was performed by treatment and sex. For goblet cell number and size, a three-way ANOVA was performed by treatment, region, and sex. All data are presented as means +/− standard error of the mean (SEM). In all instances where p values are noted, a post hoc Tukey’s multiple comparison test was performed to compare individual means. A Brown-Forsythe equality of variance test was performed for all two-way ANOVAs to confirm variances were not significantly different across groups. For three-way ANOVAs, a Spearman’s test for heteroscedasticity was used to confirm equal error.

## 3 RESULTS

### 3.1 Rotenone Treatment Increased Myenteric Plexus aSyn

To verify that mice responded to rotenone treatment, body weight was measured daily throughout the entirety of the experiment. Mice had similar starting body weights regardless of treatment group (vehicle: 24.2 g +/−5, rotenone: 21.8 g +/− 0.8) (**Fig.1A**). Change in weight was measured by subtracting starting weight from each daily weight measurement. Rotenone treatment resulted in blunted weight gain in males (**Fig. 1B**) and females (**Fig. 1C**). As a further measurement of rotenone impact, a researcher blind to treatment correctly identified treatment groups based on a five-minute open field behavior test. Rotenone treated mice exhibited decreased exploratory movement and alterations in gait, similar to our previous findings.^39^

**Figure 1.**
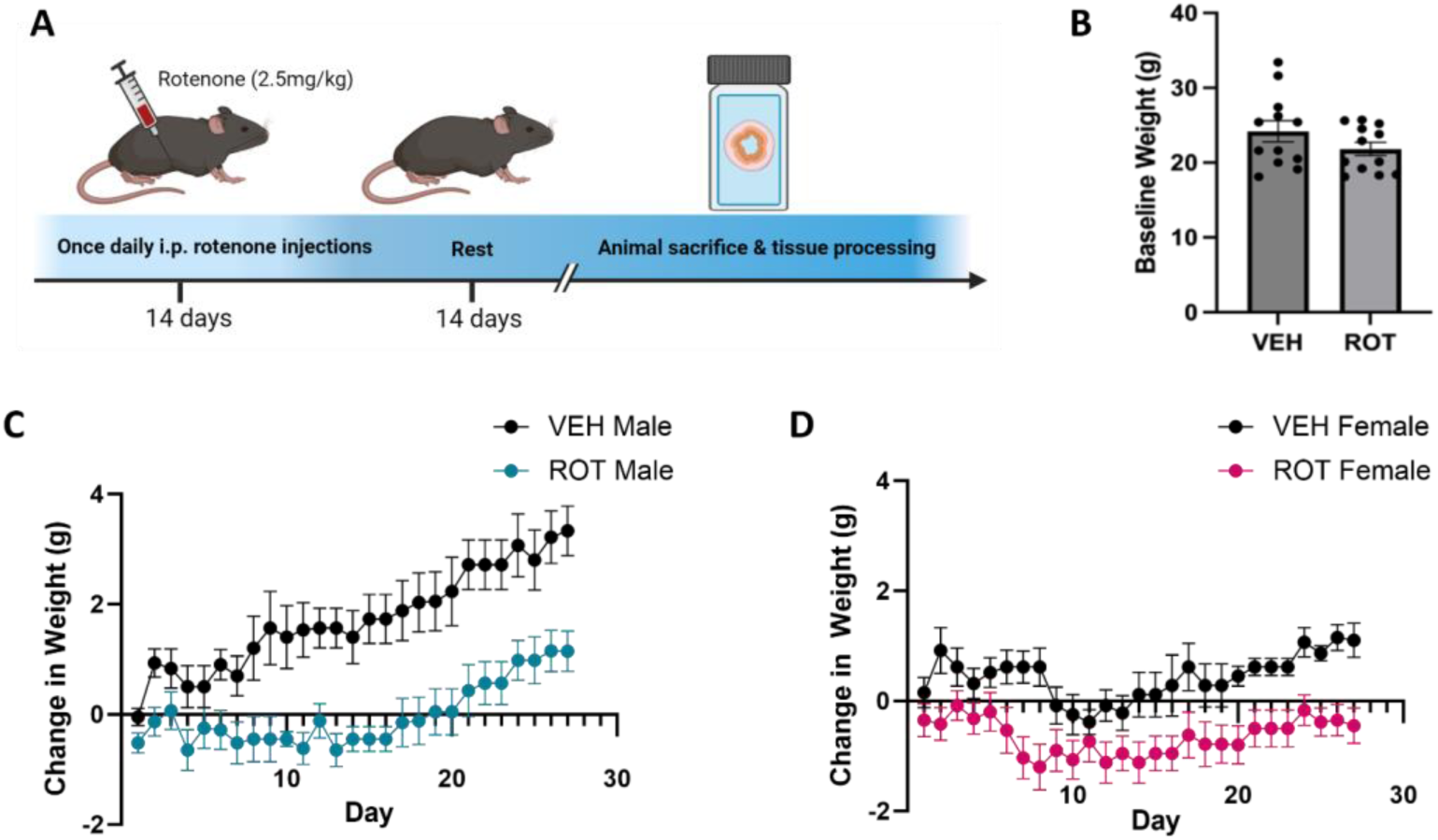
Rotenone blunted weight gain in males and females. (**A**) Overview of experimental timeline. (**B**) Baseline weights of male and female vehicle vs. rotenone mice. (**C**) Average change in weight from baseline in males. (**D**) Average change in weight from baseline in females. VEH = vehicle, ROT = rotenone.

Following behavioral confirmation of impairment, intestinal sections were analyzed for myenteric neuronal changes using a peripheral neuronal marker, PGP 9.5. Total aSyn area increased in myenteric PGP 9.5^+^ neurons in males and females after 2 weeks of rotenone treatment, with an increase in immunoreactive (ir) aSyn of 203% in females (1.5 µm^2^ +/− 0.1, p<0.0001) and 50% in males (1.3 µm^2^ +/− 0.04, p<0.05) (**Fig. 1B**) compared to vehicle treated animals (females: 0.49 +/− 0.10, p > 0.05; males: 0.84 +/−0.11, p > 0.05). A two-way ANOVA revealed a main effect of rotenone treatment (F(1,20) = 60.50, p < 0.0001) and a significant interaction between sex and rotenone (F(1,20) = 10.55, p < 0.01) indicating that rotenone’s impact was reliably greater in females. This was due in part to vehicle treated females starting with 58% (0.49 µm^2^ +/− 0.07) less immunoreactive myenteric aSyn than vehicle treated males (p<0.05, **Fig. 2B**). PGP 9.5-ir area was not different in males vs. females after rotenone treatment (Interaction effect: F(1,17) = 0.177, p > 0.05) (**Fig. 2C**) indicating that increases in aSyn area were not a result of elevated ir-PGP 9.5^+^ cell area.

**Figure 2.**
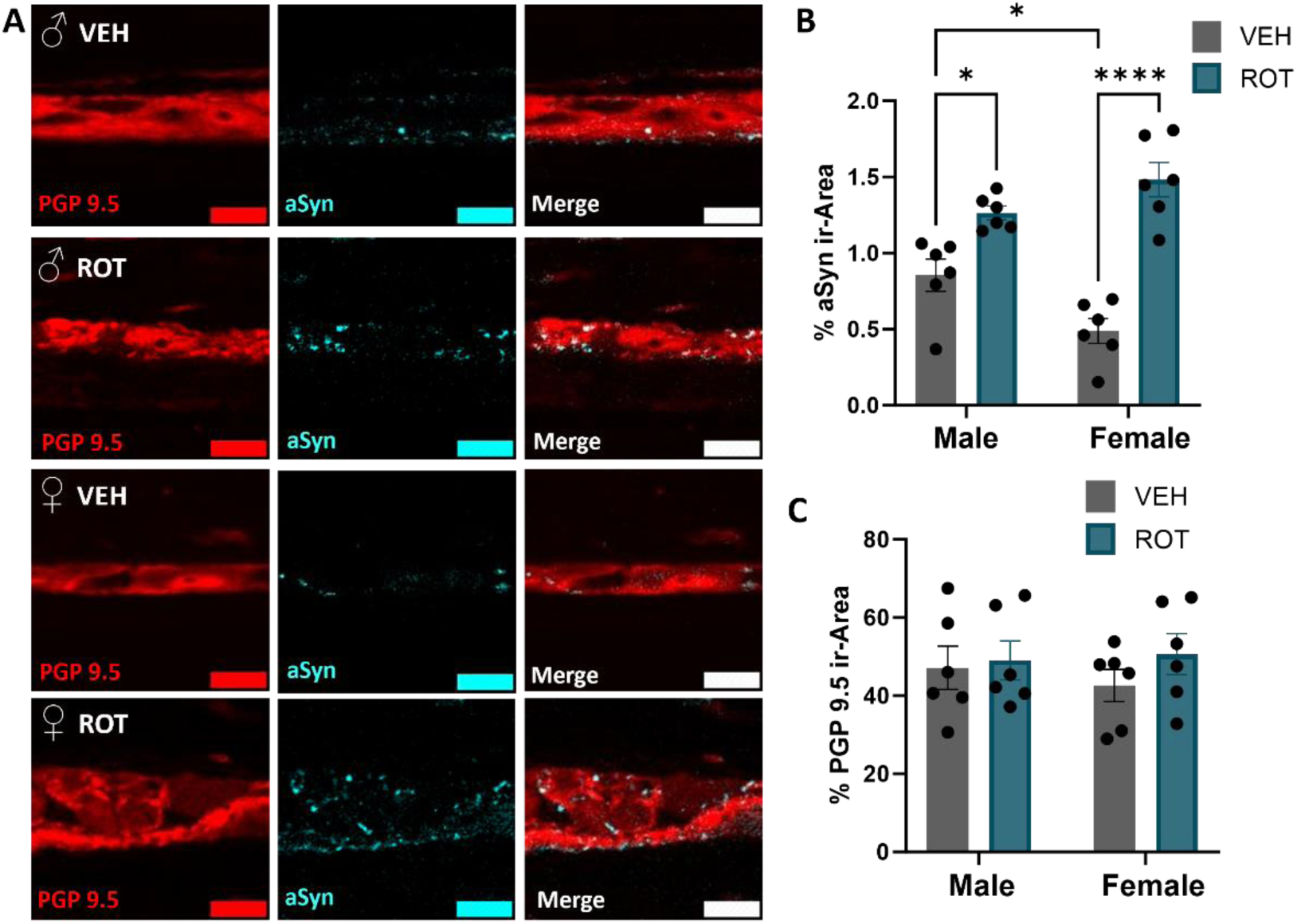
Myenteric plexus aSyn was greater in colons from rotenone treated mice. (**A**) Compared to vehicle both male and female proximal colon showed greater aSyn immunoreactivity in myenteric neurons with rotenone treatment. Females exhibited larger a larger increase in myenteric aSyn than males. (**B**) Quantification of aSyn immunoreactivity in the myenteric plexus. (**C**) Quantification of PGP 9.5 immunoreactivity in the myenteric plexus. Scale bars are 15 μm, *p<0.05, **p<0.003 ****p<0.0001. VEH = vehicle, ROT = rotneone, aSyn = alpha synuclein, PGP9.5 = protein gene product 9.5.

### 3.2 Apical CGRP Fiber Quantities Elevated in Rotenone Treated Mice

CGRP fiber immunoreactivity was interrogated by crypt region (i.e., apical crypt vs. basal crypt) in separate two-way ANOVAs. In apical crypts, CGRP levels were greater in rotenone treated animals (F(1,20) = 61.93, p < 0.0001) (**Fig. 3A,C**). This increase was 26% higher in females compared to males (F(1,20) = 9.74, p < 0.01) (**Fig. 3C**), driven by the rotenone treated groups, as there was no difference by sex in vehicle (F(1,20) = 2.89, p > 0.05) and no sex by treatment interaction.

**Figure 3.**
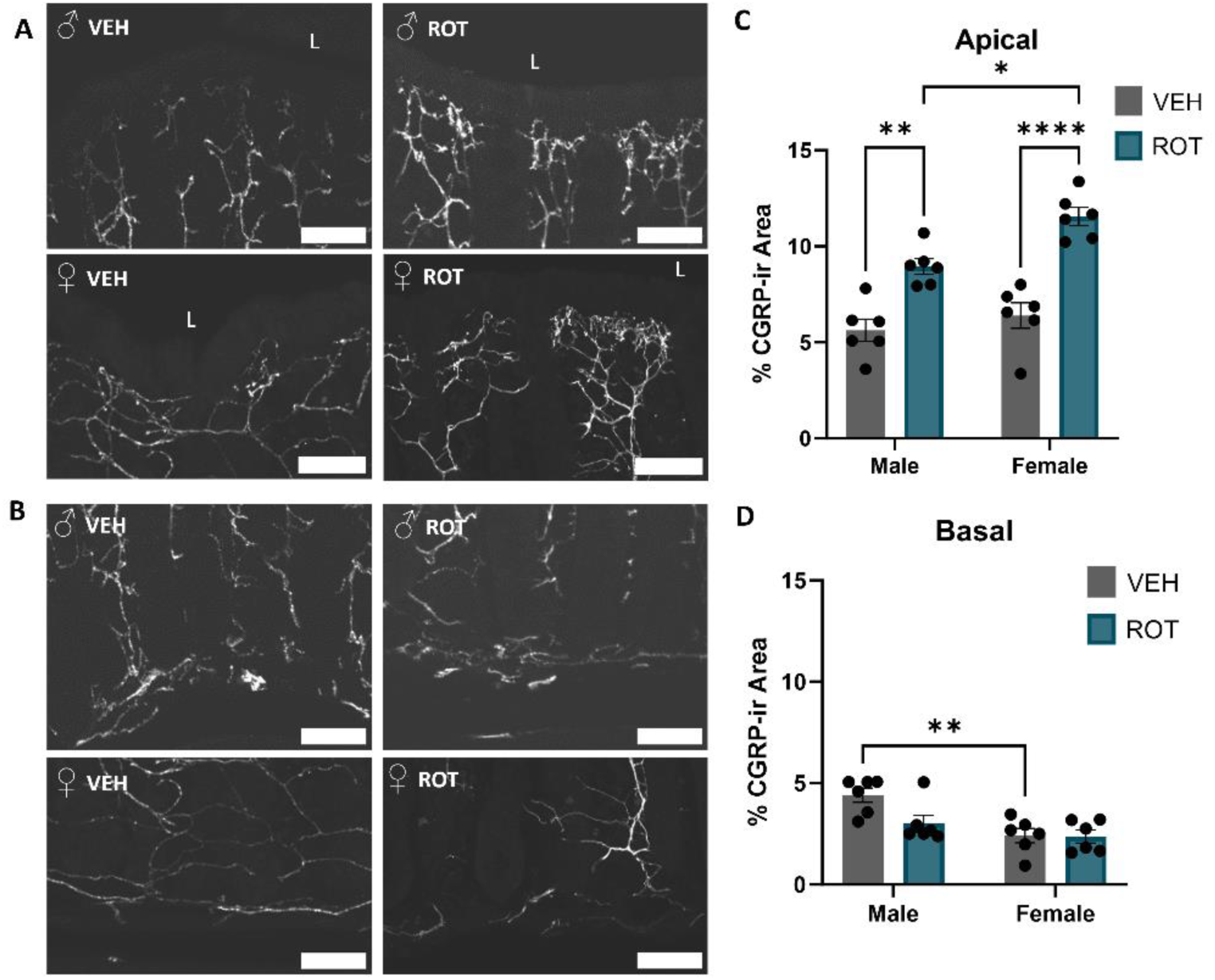
Apical CGRP^+^ fiber quantities were elevated in colon crypts from rotenone treated mice. (**A**) Rotenone treatment increased the apical percent area of CGRP^+^ fibers in males and females. (**B**) Rotenone treatment did not alter basal percent area of CGRP^+^ fibers in males and females. (**C**) Quantification of average perecent area of CGRP immunoreactivity in the apical crypt. (**D**) Quantification of average perecent area of CGRP immunoreacity in the basal crypt. Scale bars are 50 μm, L = lumen, *p<0.05, **p<0.003 ****p<0.0001. VEH = vehicle, ROT = rotenone, CGRP = calcitonin gene related peptide.

In basal crypt regions, the average percent area of CGRP-ir was low in males and females. There was, however, a main effect of sex (F(1,20) = 13.2, p < 0.05) where male vehicle treated animals had elevated basal CGRP-ir compared to vehicle treated females (**Fig. 3D**). The interaction of sex by treatment was not statistically significant suggesting some reliability to the sex effect even though the sex difference between treated mice was small.

### 3.3 Rotenone Treatment Increased aSyn in CGRP^+^ Myenteric Neurons and Apical Fibers

Immunoreactive aSyn was colocalized with CGRP^+^ myenteric neurons and apical mucosal fibers in a sex selective manner (**Fig. 4**). Two-way ANOVA investigating aSyn in CGRP^+^ myenteric neurons revealed a main effect of rotenone treatment (F(1,20) = 58.37, p < 0.0001). The percent area of aSyn in myenteric CGRP neurons (**Fig. 4A,B**) was greater in males and females treated with rotenone (male 12% +/− 1.2; female 10% +/−0.6) compared to vehicle (male 4.8% +/− 0.7, p < 0.0001; female 4.7% +/− 0.7, p < 0.005) (**Fig. 4A**). Levels of aSyn in apical CGRP^+^ fibers (**Fig. 4C,D,E**) were increased in rotenone treated mice (F(1,20) = 116.7, p < 0.0001), of both sexes (F(1,20) = 6.21, p < 0.05), however, females had a larger increase as demonstrated by a statistically significant interaction between sex and treatment (F(1,20) = 6.57, p < 0.05). The percent area of aSyn in apical CGRP fibers was elevated in males (vehicle: 0.04% +/− 0.02, rotenone: 0.9% +/− 0.1, p < 0.0001) and females (vehicle: female 0.04% +/− 0.02, rotenone: 1.5% +/− 0.2, p < 0.0001) (**Fig. 4E**). This increase was 66.57% higher in female animals compared to males (p < 0.01) (**Fig. 4E**). To confirm this increase was not due to greater apical CGRP^+^-IR in rotenone treated animals, the ratio of aSyn-IR to apical CGRP was assessed. The ratio of aSyn-IR to apical CGRP-IR (**Fig. 4F**) was increased in rotenone treated mice (F(1,20) = 47.94, p < 0.0001), of both sexes (F(1,20) = 4.88, p = 0.05), however, females had a larger increase as demonstrated by a statistically significant interaction between sex and treatment (F(1,20) = 4.879, p < 0.05). The ratio of aSyn-IR to CGRP-IR was elevated in males (vehicle: 0.0002 +/− 0.0001, rotenone: 0.003 +/− 0.007) and females (vehicle: 9.8e-05 +/− 5e-05, rotenone: 0.006 +/− 0.001) in rotenone treated mice compared to vehicle. This increase was 50% higher in female animals compared to males (p < 0.05).

**Figure 4.**
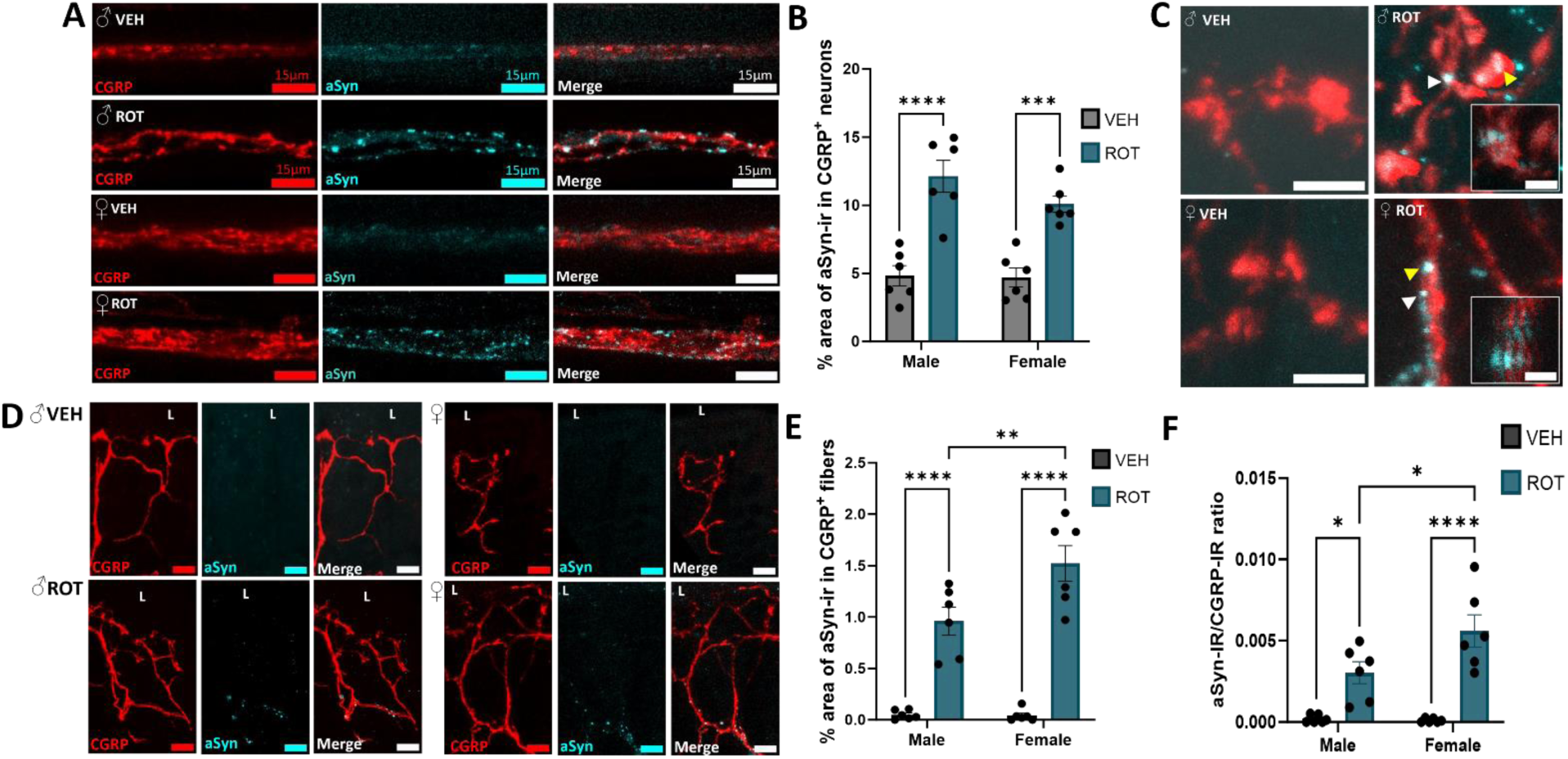
Rotenone treatment increased aSyn in CGRP+ myenteric neurons and apical fibers. (**A**) Rotenone treatement increased the percent area of aSyn in myenteric CGRP neurons in males and females. (**B**) Rotenone treatment increased the percent area of aSyn in apical CGRP fibers in males and females. (**C**) High resolution images of aSyn-ir in apcial CGRP varicosities. Arrows indicate areas of CGRP and aSyn colocalization, white arrows indicate region of inset. Insets are volume rendered cross-sections taken at a 60 degree angle. (**D**) Quantification of percent area of aSyn immunoreactivy in myenteric CGRP^+^ neurons. (**E**) Quantification of percent area of aSyn immunoreactivity in apical CGRP^+^ fibers. (**F**) Quantification of ratio of aSyn-IR in CGRP-IR fibers. Scale bars are 15 μm for A & D, for C scale bars are 5 μm and 2 μm for insets. L = lumen, * p<0.05, **p<0.01, ****p<0.0001. VEH = vehicle, ROT = rotenone, CGRP = calcitonin gene related peptide, aSyn = alpha synuclein.

### 3.4 Goblet Cells Altered Morphologically After Rotenone Treatment

Mucopolysaccharides were fluorescently labeled with the lectin Ulex Europaeus Agglutinin I (UEA1) and goblet cell size and number were analyzed by crypt region (i.e., apical vs. basal, **Fig. 5**). A three-way ANOVA revealed a main effect of region (F(1,40) = 82.41, p < 0.001) where apical goblet cell numbers were lower in animals treated with rotenone (F(1,40) = 12.92, p < 0.001) regardless of sex (F(1,40) = 1.4, p > 0.05) (**Fig. 5A-C**). Males treated with rotenone had 53% fewer apical goblet cells (5.0 +/− 0.1) (**Fig. 5B**) compared to vehicle (10.0 +/− 0.4, p < 0.0001) (**Fig. 5A, C**) and rotenone treated females had 48% fewer cells (4.0 +/− 0.2) compared to vehicle (8.0 +/− 0.5, p < 0.0001). Vehicle treated males had 20% higher baseline goblet cell counts compared to females restricted to the apical region (F(1,40) = 13.07, p < 0.001) (**Fig. 5C**). There were 20% fewer basal goblet cells after rotenone treatment in males (10.0 +/− 0.6) (**Fig. 5B’**) compared to vehicle (8.0 +/− 0.3, p < 0.05) (**Fig. 5A’**) that did not reach statistical significance by posthoc testing in females (**Fig. 5C**).

**Figure 5.**
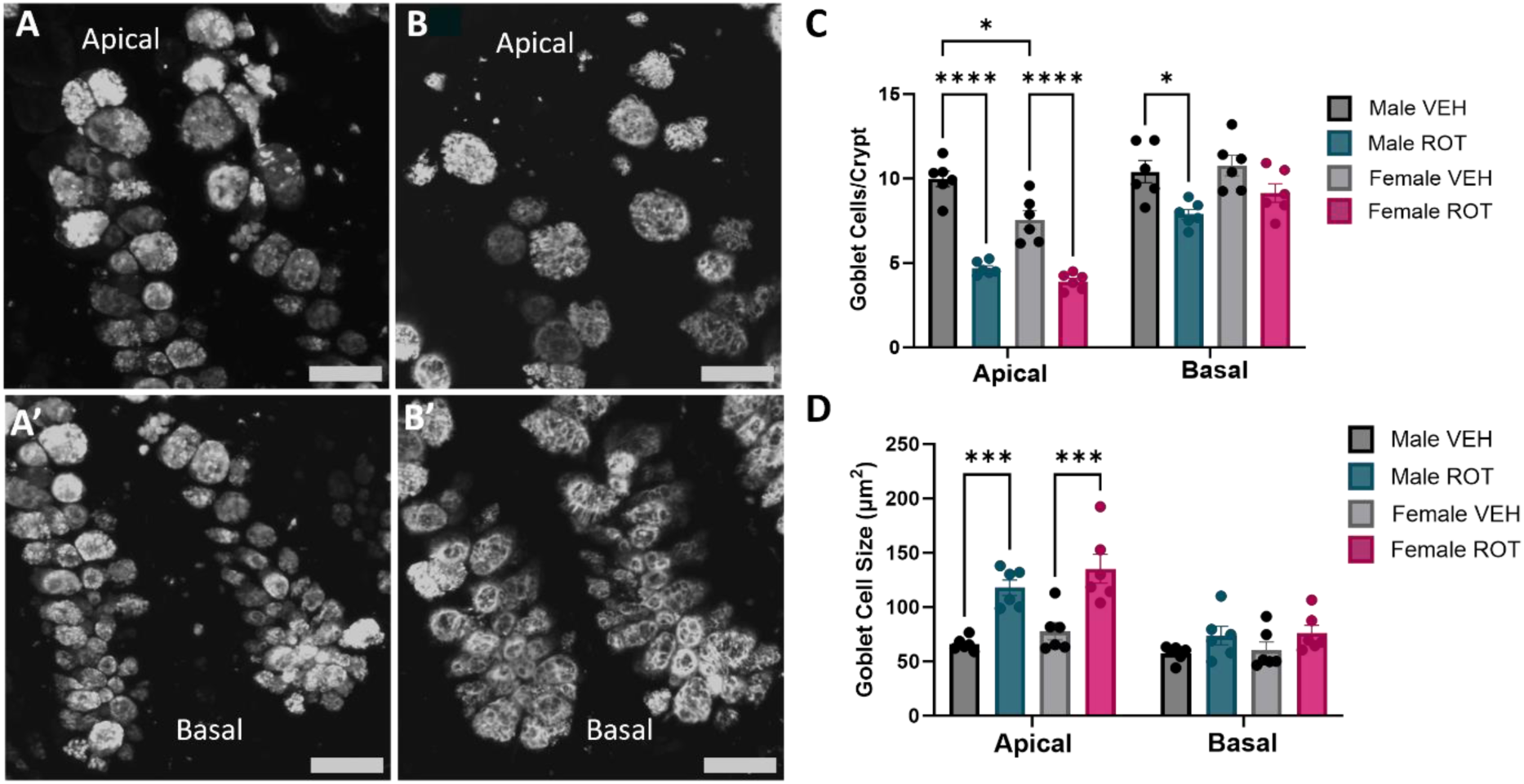
UEA1^+^ goblet cells altered morphologically after rotenone treatment. (**A**) UEA1^+^ goblet cells at the apex and (**A’**) base of the crypt in vehicle treated tissue have a smaller average immunoreactive area and are fewer in number than (**B**) UEA1^+^ goblet cells at the apex and (**B’**) base of the crypt in rotenone treated tissue. (**C**) Quantification of UEA1^+^ goblet cell number and (**D**) UEA1^+^ immunoreactive area. Scale bars 25 μm, representative images are male, *p<0.05, ***p<0.001 ****p<0.0001. VEH = vehicle, ROT = rotenone, UEA1 = Ulex Europaeus Agglutinin I

In contrast to fewer goblet cells after rotenone, those goblet cells that remained were larger. A three-way ANOVA investigating goblet cell size revealed a main effect of region (F(1,40) = 34.34, p < 0.001) where goblet cell size was greater in rotenone treated animals selectively in the apical crypt regions (F(1,40) = 12.36, p < 0.001), with no main effect of sex (F(1,40) = 1.14, p > 0.05) (**Fig. 5D**). There was a main effect of treatment (F(1,40) = 40.95, p < 0.0001) where apical goblet cell size was 79% greater in rotenone treated males (117.7 µm^2^ +/ 6.1) (**Fig. 5B**) compared to vehicle treated males (65.9 µm^2^ +/− 2.1, p < 0.001) (**Fig. 5A**) and 74% greater in rotenone treated females (135.3 µm^2^ +/− 11.2) compared to those treated with vehicle (77.8 µm^2^ +/− 6.7, p < 0.001).

## 4 DISCUSSION

Females with PD exhibit faster disease progression and higher mortality rates than males^10^. Using a rotenone mouse model of PD, the results of the current study point towards sex selective changes in enteric aSyn levels, alterations in CGRP distribution, and changes in goblet cell morphology, potentially influencing the more severe outcomes in females with PD. These findings contribute to our understanding of the link between intestinal homeostasis and aSyn aggregation in peripheral PD pathology and may provide insights for advancing early diagnostic strategies. Few studies of peripheral aSyn aggregation or PD pathological etiology include sex as a biological variable. Given that females with PD exhibit a higher mortality rate and report increased severity of GI symptoms,^10,11,43^ the present studies provide an indication of potential sex-selective targets of a disease process.

Enteric aSyn levels are elevated in PD,^44^ and the pesticide rotenone increases aSyn in the central nervous system^39^ and the GI tract.^45^ The present study demonstrated that intraperitoneal rotenone injections increased total aSyn in myenteric plexus neurons consistent with pathology found after oral rotenone administration.^45^ There was a basal sex difference in myenteric aSyn levels, with males having higher quantities. When treated with rotenone, aSyn levels rose significantly in both sexes, but more so in females. This finding potentially implicates the female myenteric neuronal population as having an altered neurochemical phenotype at baseline, akin to the estrus cycle dependent change in neuronal nitric oxide synthase (nNOS) neuron phenotype.^46^ The lower baseline but subsequent post-rotenone compensation of aSyn levels in female CGRP neurons and fibers may influence GI motility and goblet cell secretory function. Together, these data provide a potential explanation for the more severe GI transit dysfunction in PD females. Further investigations of the impact of aSyn on GI motility and neuroepithelial signaling within the context of sex are clearly warranted.^11,14^

Neurons utilizing the neuropeptide CGRP for signaling may be critically involved in regulating intestinal motility and nociception.^31,47,48^ Visceral pain is regulated by gut wall components, including the secretions from the microbiome adjacent to the colonic wall, and via enteric neuronal afferent CGRP signaling to the dorsal root ganglia.^22,35,48^ Further, agonizing colonic CGRP receptors prior to colonic distention yields elevated visceral pain responses in conventionally colonized mice^22^. The increased CGRP^+^ apical fiber density observed in this study may contribute to elevated sensitivity to noxious stimuli from the lumen, however, receptor expression patterns for noxious ligands on apical CGRP^+^ fibers were not investigated in this study. The greater increase in apical CGRP in rotenone-treated females provides a focal point highlighting CGRPs potential involvement in the higher incidence of visceral pain observed in female PD patients. Studies examining CGRP receptor populations along with peptide release in these anatomically distinct enteric regions are needed to identify functional connections among the gut circuits identified here.

Alpha-synuclein aggregation can elicit morphological and firing pattern changes in neuronal populations.^49^ Increased aSyn in CGRP^+^ enteric neurons may modulate downstream CGRP anatomical alterations and signaling patterns. CGRP production is upregulated in damaged nerves^25^ and intraperitoneal injections of the CGRP antagonist BIBN 4096 BS decreased central aSyn expression in a mouse model of Alzheimer’s disease.^50^ Rotenone treated males and females had increased aSyn in CGRP^+^ myenteric neurons and apical CGRP^+^ fibers, suggesting that aSyn levels may be related to alterations in gut CGRP neuronal immunoreactivity akin to those seen in PD patient derived neuronal cultures.^49,51^ Female mice showed a greater increase in aSyn in apical CGRP^+^ fibers, a greater increase in immunoreactive CGRP in apical regions of colonic crypts, and lower baseline levels of myenteric aSyn compared to males suggesting that females might be more susceptible to aSyn and CGRP localization within neuronal fibers following rotenone treatment.

Underlying causes of constipation, one of the earliest symptoms of PD,^15,16^ are multifaceted and treatment with anti-CGRP monoclonal antibodies decreases GI transit.^31,32^ Similarly, animal studies with rotenone report decreased intestinal transit.^8,9^ In the present study, rotenone treated mice exhibited increased area of CGRP immunoreactive fibers at the apical crypt. One explanation for the association between elevated gut mucosal CGRP and potential aSyn driven constipation is through regulation of mucus production by goblet cells. In the current study there were fewer goblet cells that were larger following rotenone treatment. The impact was stronger apically, coinciding with the greater CGRP fiber density after rotenone. Goblet cells originate from progenitor cells at the base of the crypts and mature as they migrate apically where they acquire the ability to synthesize and secrete mucoplysaccharides.^52^ Mature goblet cells are located in the upper third of crypts^53,54^ potentially making them more susceptible to morphological changes due to local environmental factors like dense CGRP producing neuronal fibers. While acute stimulation of goblet cells serves as a protective mechanism to bolster the mucosal barrier in response to pathogens,^55^ chronic stimulation may lead to goblet cell mucus depletion. Impacts of aSyn driven alterations to goblet cell mucus production and secretion remains unclear. The current results suggest a need to investigate goblet cell mucus production and mucosal thickness to determine if altered CGRP signaling after rotenone leads to mucosal barrier dysfunction.

## 5 CONCLUSIONS

In conclusion, rotenone treatment reproduced PD clinical findings of increased aSyn in the myenteric plexus and extended these findings to show increases in apical CGRP immunoreactivity and altered goblet cell morphology in colonic crypts. Enteric aSyn may alter CGRP neuronal interactions with goblet cells that are critical for mucus production or secretion. These findings demonstrate a potential pathway for sex selective neuronal-epithelial interaction between CGRP and goblet cells secondary to elevated enteric aSyn, identifying a potential contributor to GI dysfunction in PD. Since GI symptoms arise in a sex specific manner, years before motor dysfunction, understanding mechanisms of enteric aSyn pathophysiology is crucial for identifying early diagnostic markers and novel therapeutic targets.

## ACKNOWLEDGMENTS

We would like to thank Connie King^1^ for help with animal processing and Casey P McDermott^2^ for help with initial trial experiments and rotenone dosing protocol. We would also like to acknowledge the College of Veterinary Medicine and Biomedical Sciences Graduate Research Scholar Award for their partial support of these studies.

^1^ Department of Biomedical Sciences, Colorado State University, Fort Collins, CO, USA

^2^ Department of Environmental and Radiological Health Sciences, Colorado State University, Fort Collins, CO, USA

## AUTHOR CONTRIBUTIONS

**Hayley N Templeton:** Conceptualization, Formal Analysis, Investigation, Writing-Original Draft, Visualization. **Alexis T Ehrlich:** Investigation, Writing-Review & Editing. **Luke A Schwerdtfeger:** Conceptualization, Formal Analysis, Writing-Review & Editing, Supervision. **Julietta A Sheng:** Investigation. **Ronald B Tjalkens:** Methodology, Resources**. Stuart A Tobet:** Conceptualization, Supervision, Project Administration, Funding Acquisition, Writing-Review & Editing.

## Supplemental figures

**Supplemental Figure 1.**
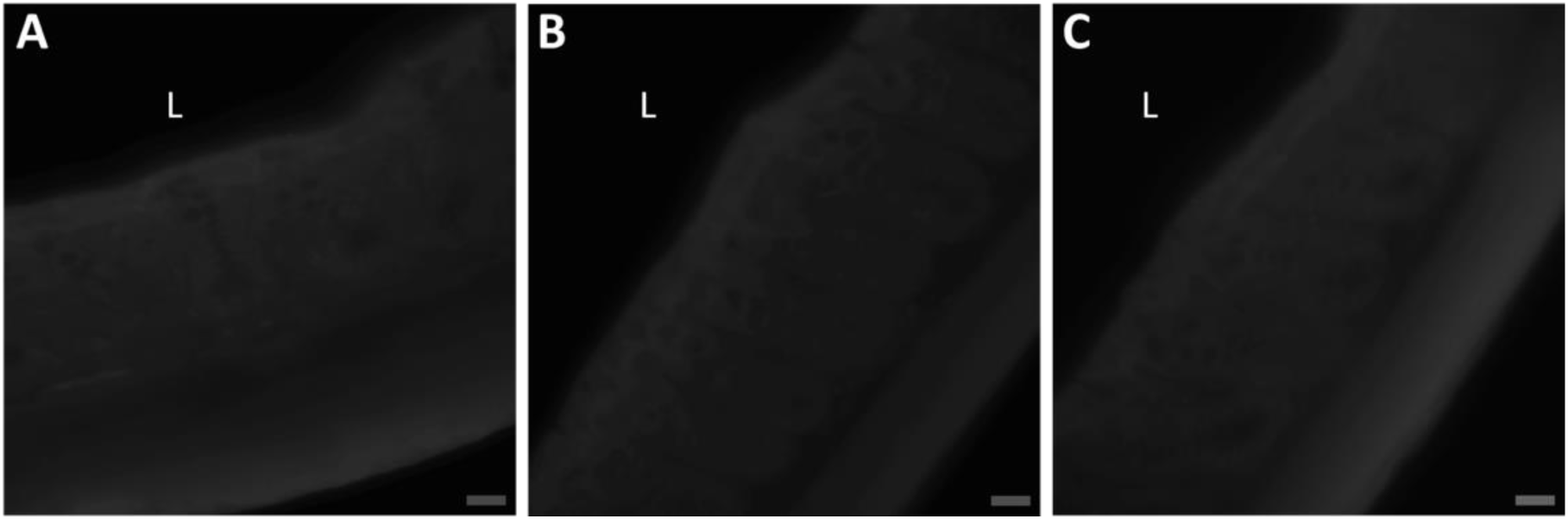
CGRP and alpha synculein antibody controls. (**A**) Pre-absorption with αCGRP peptide. (**B**) αCGRP negative control. (**C**) aSyn negative control. Scale bars are 50 µm, L= lumen.

